# The endogenous *Mtv8* locus and the immunoglobulin repertoire

**DOI:** 10.1101/2023.11.24.567955

**Authors:** Helen A. Beilinson, Steven A. Erickson, Tatyana Golovkina

## Abstract

The vast diversity of mammalian adaptive antigen receptors allows for robust and efficient immune responses against a wide number of pathogens. The antigen receptor repertoire is built during the recombination and hypermutation of B and T cell receptor (BCR, TCR) loci. V(D)J recombination rearranges these antigen receptor loci, which are organized as an array of separate V, (D), and J gene segments. Transcription activation at the recombining locus leads to changes in the local three-dimensional architecture, which subsequently contributes to which gene segments are utilized for recombination. The endogenous retrovirus (ERV) mouse mammary tumor provirus 8 (*Mtv8*) resides on mouse chromosome 6 interposed within the large array of light chain kappa V gene segments. As ERVs contribute to changes in genomic architecture by driving high levels of transcription of neighboring genes, it was suggested that *Mtv8* could influence the BCR repertoire. We generated *Mtv8*-deficient mice to determine if the ERV influences V(D)J recombination to test this possibility. We find that *Mtv8* does not influence the BCR repertoire.

## Introduction

The extensive diversity of the jawed vertebrate adaptive immune response depends on the programmed assembly and hypermutation of antigen receptor (AgR) genes (*1*). The first stage of AgR assembly is V(D)J recombination, initiated by lymphocyte-specific recombination activating genes 1 and 2 (RAG1 and RAG2), during which immunoglobulin (Ig) and T cell receptor (TCR) genes are recombined from discrete variable (V), diversity (D), and joining (J) gene segments (*2*).

AgR assembly is a sequential process during lymphocyte development. In B cells, the Ig heavy-chain (*Igh*) locus recombines in early and pro B cells prior to the kappa and lambda light chain (*Igk, Igl*) loci in pre B cells (*2, 3*). *Igh* recombines in two phases: first, D_H_-to-J_H_ rearrangements occur in lymphoid progenitors; second, V_H_-to-DJ_H_ rearrangements occur in pro-B cells (*2, 4*). After successful (i.e. in-frame without premature stop codons) *Igh* recombination, *Igk* undergoes V_K_-to-J_K_ recombination (*3*). If neither *Igk* allele successfully rearranges, the *Igl* locus recombines in a V_L_-to-J_L_ fashion. During V(D)J recombination, RAG (a heterotetramer composed of RAG1 and RAG2 molecules) accumulates at recombination centers (RCs) that encompass J or DJ gene segments (for D-to-J/V-to-J and V-to-DJ recombination, respectively) (*4*). In RCs, RAG binds to a recombination signal sequences (RSS) that flanks the rearranging gene segment (*4*). Then, a partner RSS of the second gene segment is brought into the RC for synapsis and RAG-mediated cleavage.

V(D)J recombination determines the V, (D), and J gene segments used in a particular AgR gene and is dependent on three-dimensional chromosomal architecture. Specifically, V(D)J recombination is constrained to AgR loci by chromatin loops, the bases of which are defined by CCCTC-binding factor (CTCF)-bound CTCF-Binding Elements (CBEs) (*5-9*). The number of loops formed is dependent on the recombining locus (*10*). In addition to chromatin loops defining the AgR, chromosomal architecture also defines the mechanism by which recombining gene segments are brought into the RC (*11*).

The *Igk* locus is comprised of 92 functional (163 total) V_K_ and four functional J_K_ spread across 3.2 Mb (Fig. 1A) and contracts into a recombination-competent chromosomal structure in developing pro- and pre-B cells (*12, 13*). Once contracted, the *Igk* locus forms a rosette-like structure with five V_K_-containing loops and one loop with J_K_ and C_K_ gene segments (*10*). *Igk* rearrangement occurs predominantly through the collision of, and subsequent recombination between, the J_K_-containing and a V_K_-containing loop (*10*). While the *Igk* loops are thought to be predominantly shaped by the CBEs throughout the locus, it is unknown whether other factors are involved.

**Figure 1.**
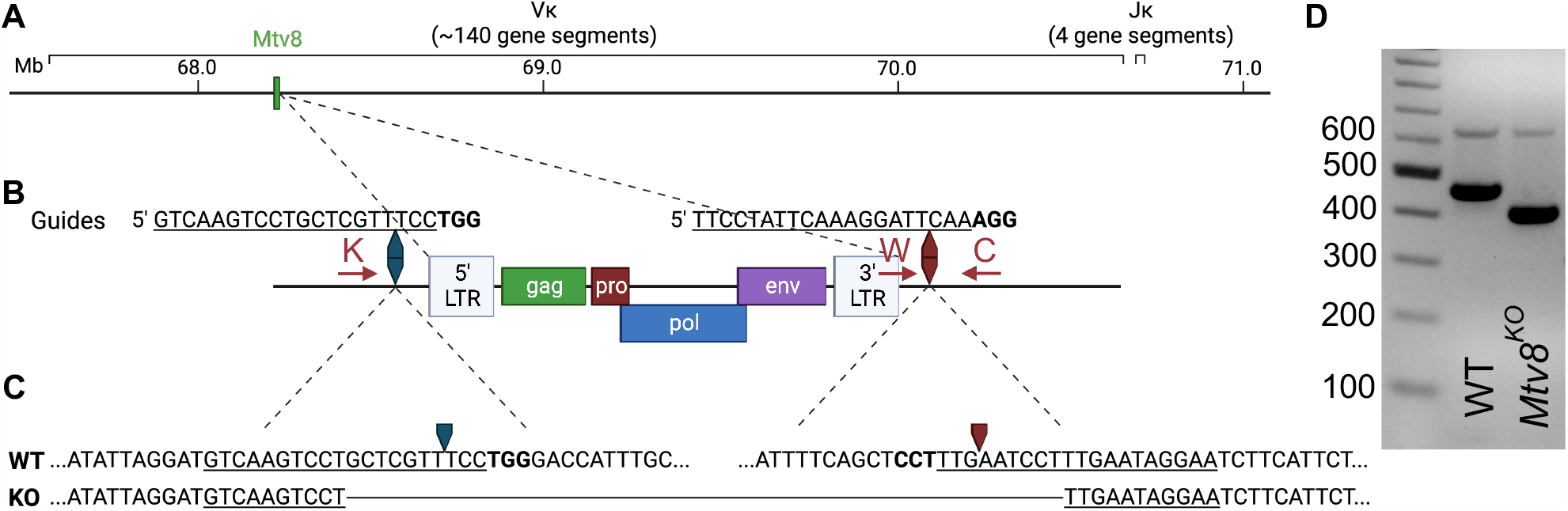
Generation of *Mtv8*^KO^ mice. (A) Schematic of B6J murine *Igk* locus on chromosome 6. Location of *Mtv8* is highlighted in green. (B) Schematic of *Mtv8* and guides used for CRISPR/Cas9 targeting of *Mtv8* to generate *Mtv8*^*KO*^ mice. LTR: long-terminal repeat; *gag*: group-specific antigen, *pro*: protease, *pol*: polymerase, *env*: envelope. Red arrows: genotyping primers (K: KOF, W: WTF, C: CommonR). (C) Genomic sequence of WT and KO *Mtv8* alleles. Only the flanking sequnces are shown for the WT allele. For (B) and (C), underlined text indicates guide sequence, bold text indicates PAM sequence, arrows indicate Cas9-cut site. (D) Gel image of genotyping representative WT and homozygous *Mtv8*^*KO*^ mice. Mice were genotyped using the KOF, WTF, and CommonR primers. WT product: 479 bp, KO product: 419 bp.

The endogenous retrovirus (ERV) mouse mammary tumor virus (MMTV), *Mtv8* was mapped to the *Igk* locus on chromosome 6 (Fig. 1A) (*14-16*). *Mtv8* is a provirus with all open-reading frames (ORFs) intact, namely, *gag, pro, pol, env*, and superantigen (SAg) (Fig. 1B). The provirus does not produce infectious virions and is silent in the mammary glands, the targeted tissue of all MMTVs, potentially due to hypermethylation of its promoter region (*17-19*).However, expression of the provirus is detectable in lymphoid cells (*20*). As *Mtv8* is mapped within the *Igk* locus, it was hypothesized that it may contribute to *Igk* recombination by driving high levels of transcription in its vicinity (*21*). Previously, to identify whether *Mtv8* affects the *Igk* repertoire, the frequency of recombination of the J_K_ gene segments to the first V_K_ gene segment downstream of *Mtv8* (V_K_14-111, formerly V_K_9M) was analyzed in inbred mouse strains with (BALB/c, C58.C, A/J, and C57BL/6J (B6J)) and without (C58, C.C58, NZB, and PERA/Ei) *Mtv8 (21*). These experiments demonstrated that mice inheriting *Mtv8* have higher recombination between V_K_14-111 and all J_K_ gene segments compared to strains without *Mtv8*. While these analyses suggested differences in the usage of V_K_14-111, *Ig* repertoire comparisons between inbred mouse lines is not an ideal approach, as polymorphisms within the loci other than *Mtv8* could also influence the repertoires.

To address whether *Mtv8* shapes the repertoire of mice with this ERV, we used CRISPR/Cas9 technology to generated B6J mice lacking *Mtv8* and compared their *Ig* repertoire to that of wild-type (WT) B6J mice. We found that the absence of *Mtv8* had no significant effects on the *Igh, Igk*, and *Igl* gene segment usage. Thus, contribution of *Mtv8* to the mouse *Ig* repertoires can be definitively ruled out.

## Results and Discussion

### Generation of Mtv8-deficient B6J mice

*Mtv8* knockout (KO) B6J mice were generated using a CRISPR/Cas9 approach. To target *Mtv8* without disturbing the V_K_ gene segments in its vicinity, we designed two guides to precisely delete the ERV: one 645 bp upstream of the 5’ LTR and one 79 bp downstream of the 3’ LTR (Fig. 1B). A founder was identified using PCR with primers flanking the predicted deleted region and it was determined that a 10,688 bp resection occurred, resulting in the deletion of the entire *Mtv8* locus (Fig. 1C, D). The founder was crossed to a wildtype (WT) B6J mouse and heterozygous, mutant allele-carrying progeny were interbred to generate a homozygous *Mtv8*-deficient line (Fig. 1D).

### Mtv8 does not contribute to light and heavy chain recombination

To investigate whether *Mtv8* alters *Ig* gene segment usage, we analyzed the Ig repertoires of WT and *Mtv8*^*KO*^ B6J mice (Fig. 2-4). Accordingly, RNA isolated from CD19^+^ splenocytes was used to prepare Ig heavy and light chain-specific 5’-RACE libraries, which were analyzed to determine the BCR repertoire using the pRESTO toolkit (*22*). Both the total and productive repertoire was analyze. Productive transcripts were defined as in-frame without premature stop codons and are likely those transcripts that are translated into expressed Ig proteins. The total repertoire includes transcripts that are non-translated.

**Figure 2.**
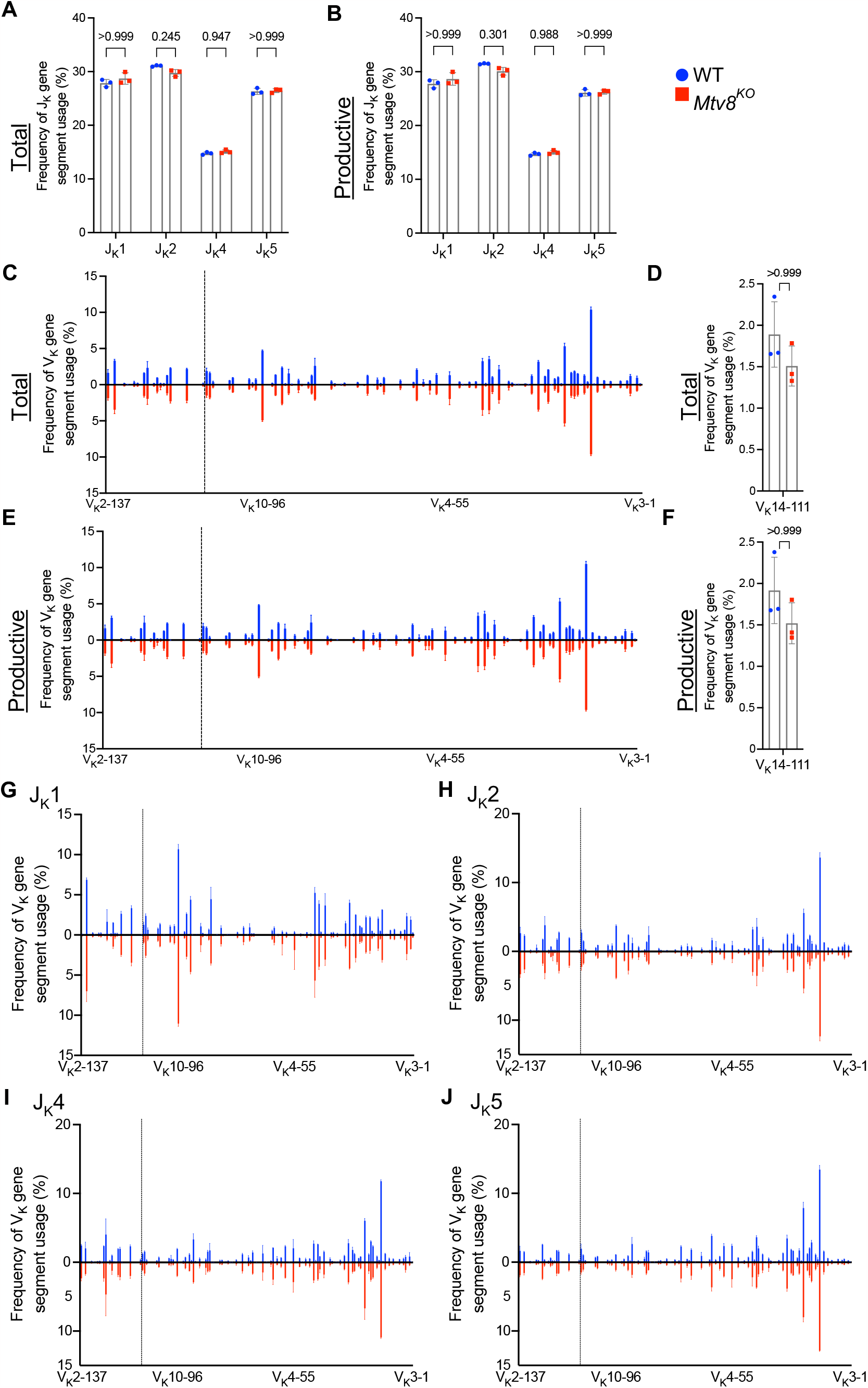
*Igk* repertoire of CD19^+^ splenocytes of *Mtv8*^*KO*^ mice. Frequency of J_K_ (A-B) and V_K_ (C-F) gene segments in total (A, C, D) and productive *Igk* (B, E, F) transcripts from CD19^+^ splenocytes. Productive recombination events are in-frame without premature stop codons. (D, F) Frequency of V_K_14-111 (previously termed VK9M) in total (D) and productive (F) *Igk* transcripts. (G-J) Frequency of V_K_ gene segments in total *Igk* transcripts recombined to J_K_1 (G), J_K_2 (H), J_K_4 (I), and J_K_5 (J). (C, E, G-J) V_K_ gene segments are arranged from 5’ distal to 3’ proximal and the vertical dotted line represents chromosomal location of *Mtv8*. n=3 mice per group. Data presented as mean with error bars indicated SD. Statistical significance was determined by unpaired Welch’s *t* test with Bonferroni correction (numbers above bars indicate adjusted P value).

To determine whether *Mtv8* influences *Igk* recombination, we examined the overall J_K_ and V_K_ usage in total and productive *Igk* transcripts from CD19^+^ splenocytes (Fig. 2). There was no difference in J_K_ usage in either total or productive transcripts (Fig. 2A, B). Furthermore, the loss of *Mtv8* led to no alterations in the total V_K_ repertoire in both total and productive recombination events (Fig. 2C, E). Notably, there was no change in usage of V_K_ gene segments mapped in the close proximity with *Mtv8* (dotted line; Fig. 2C, E). An increased usage of the V_K_ gene segment, V_K_14-111 (formerly called V_K_9M) directly downstream of *Mtv8* in *Mtv8*^*+*^ mouse strains compared to *Mtv8*^*-*^ strains was previously reported (*21*). We observed no difference in frequency of V_K_14-111 usage in total and productive *Igk* transcripts between WT and *Mtv8*^*KO*^ B6J mice (Fig. 2D, F). Thus, the previously observed disparities in V_K_14-111 usage between *Mtv8*^*+*^ and *Mtv8*^*-*^ mice from distinct genetic backgrounds are independent of *Mtv8*.

The same *Igk* locus can undergo multiple rounds of recombination. While *Mtv8* does not affect the overall frequency of V_K_ usage, we wanted to test the possibility that it might affect either the first or subsequent recombination in distinct ways. Recombination of any V_K_ to J_K_1 can only only occur during a primary recombination event, as recombination to any other J_K_ gene segment would remove J_K_1 from the *Igk* locus. Recombination to J_K_2, J_K_4, or J_K_5 can either occur during a primary or secondary recombination event. As such, we calculated the frequency of V_K_ gene segment usage in total recombination events with each J_K_ gene segment. We found that, in line with globally unaffected J_K_ gene segment usage, frequency of any particular V_K_ gene segment recombining to J_K_1, J_K_2, J_K_4, and J_K_5 was unchanged with the loss of *Mtv8* (Fig. 2E-H). Taken together, these data show that the *Igk* repertoire of *Mtv8*^*KO*^ B6J mice does not differ from WT B6J and that *Mtv8* does not influence the *Igk* recombination.

We also found no difference in the J_L_ and V_L_ usage among total and productive *Igl* transcripts between WT and *Mtv8*^*KO*^ B6J mice (Fig. 3). Similarly, we found no alterations in J_H_, D_H_, and V_H_, and gene segment between WT and *Mtv8*^*K*O^ mice (Fig. 4).

**Figure 3.**
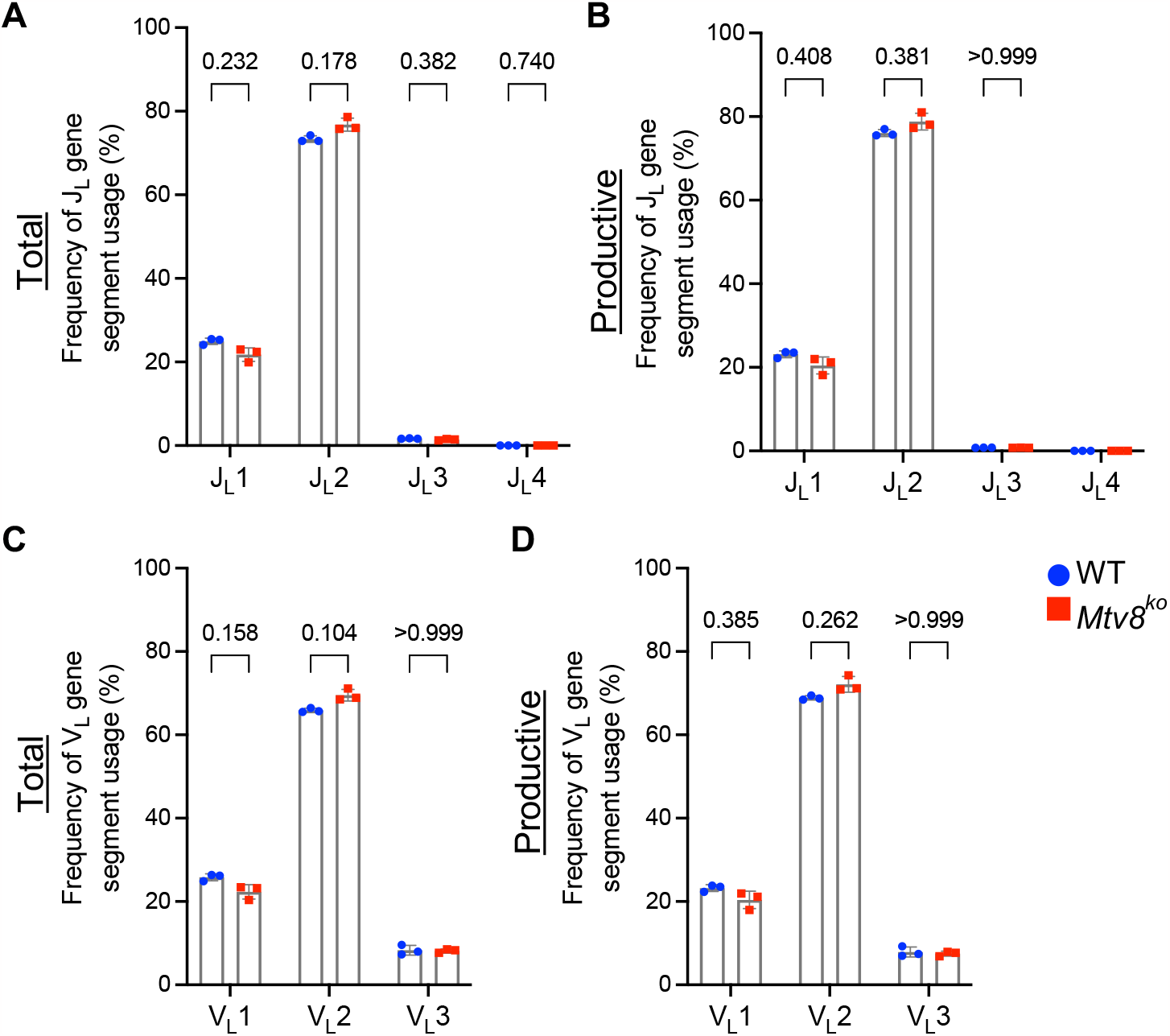
*Igl* repertoire of CD19^+^ splenocytes of *Mtv8*^*KO*^ mice. Frequency of J_L_ (A-B) and V_L_ (C-D) gene segments in total (A, C) and productive (B, D) *Igl* transcripts from CD19^+^ splenocytes. Productive recombination events are in-frame without premature stop codons. n=3 mice per group. Data presented as mean with error bars indicated SD. Statistical significance was determined by unpaired Welch’s *t* test with Bonferroni correction (numbers above bars indicate adjusted P value).

**Figure 4.**
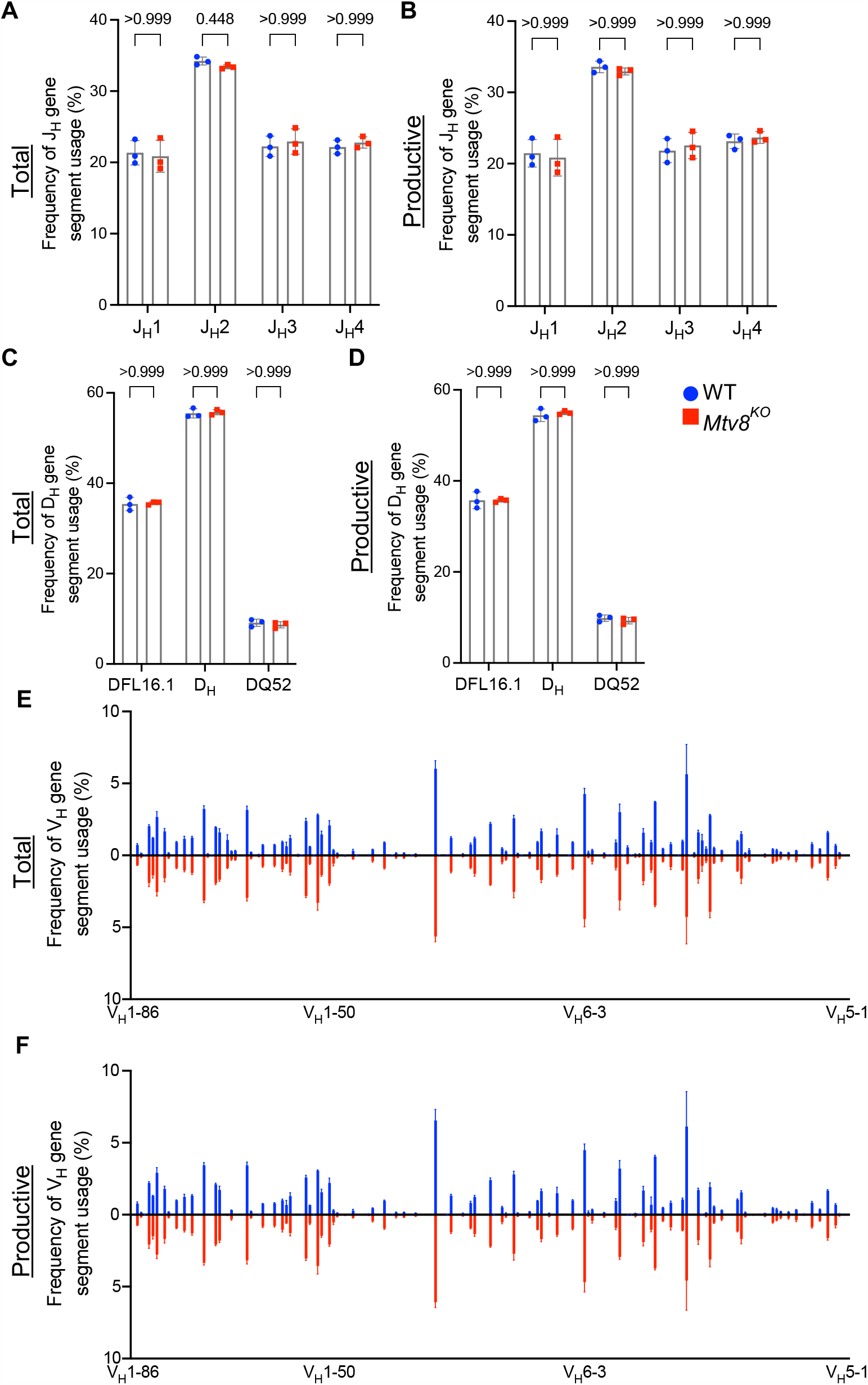
*Igh* repertoire of CD19^+^ splenocytes of *Mtv8*^*KO*^ mice. Frequency of J_H_ (A-B), D_H_ (C-D), and V_H_ (E-F) gene segments in total (A, C, E) and productive (B, D, F) *Igh* transcripts from CD19^+^ splenocytes. Productive recombination events are in-frame without premature stop codons. (C-D) DSP gene segments are the intervening D_H_ gene segments between DFL16.1 and DQ52 and were considered as a group (*24*). (E-F) V_H_ gene segments are arranged from 5’ distal to 3’ proximal. (G-H) Frequency of V_H_ gene segments in total (G) and productive (H) *Igh* transcripts that are statistically different between WT and *Mtv8*^KO^ mice. V_H_ gene segments are arranged from 5’ distal to 3’ proximal. n=3 mice per group. Data presented as mean with error bars indicated SD. Statistical significance was determined by unpaired Welch’s *t* test with Bonferroni correction (numbers above bars indicate adjusted P value).

It is now accepted that the collisions of the rosette-like loops of the *Igk* locus are the major mechanism of V_K_-to-J_K_ recombination (*10*). However, the identification of an ERV, *Mtv8*, in the middle of the V_K_ array on the *Igk* locus led to the hypothesis that it may influence the Ig repertoire (*14-16*). Differences in V_K_ usage in mouse strains with and without *Mtv8* initially supported this hypothesis (*21*). Now, advancements in genome-editing technologies allow us to definitely address whether a genetic loss of *Mtv8* can change the *Ig* repertoire within a mouse line. We show that *Mtv8* has no effect on the *Ig* repertoire by analyzing the *Igh, Igk*, and *Igl* transcriptional repertoires of B cells isolated from WT and *Mtv8*^KO^ B6J mice. These data demonstrate that while *Mtv8* is an intact ERV at the *Igk* locus, transcription driven by its LTRs has no influence on the shaping of the *Igk* loops, and thus, *Igk* recombination.

## Methods

### Mice

Mice utilized in this study were bred and maintained at the animal facility of The University of Chicago. The studies described herein have been reviewed and approved by the Animal Care and Use Committee at the University of Chicago, which is accredited by the Association for Assessment and Accreditation of Laboratory Animal Care (AAALAC International).

C57BL/6J (B6J) mice were purchased from The Jackson Laboratory. *Mtv8*^*KO*^ B6J mice were generated using CRISPR/Cas9 technology. Two guide RNAs targeting the flanking up- and downstream sequences of *Mtv8* (5’ guide: 5’-GTCAAGTCCTGCTCGTTTCC; 3’ guide: 5’-TTCCTATTCAAAGGATTCAA) were co-injected into a single cell B6J embryos along with Cas9 (Fig. 1B). Founder mice and subsequent offspring in which *Mtv8* was eliminated were identified using PCR with primers flanking the guide cut sites (KOF: 5’-GAATTTGGGTGCTCTTGCAT; CommonR: 5’-AACACAAATGGAGGCAAAGC; KO product size: 419 bp). To identify mice with the WT allele, a separate PCR was used using the CommonR primer and a WT-specific reverse primer that lays in the excited region (WTF: 5’-GGATTTGCAAACAAGATCCAG; WT product size: 479 bp). A founder line was established in which *Mtv8* was eliminated with a 10,688 bp deletion (Fig. 1C). The founder mouse was bred to a WT B6J mouse and the resulting F1 offspring were intercrossed to generate a homozygous KO line. The deletion was confirmed at the DNA level by sequencing the KO allele using the KO-specific PCR primers.

### RNA isolation from splenic B cells

CD19^+^ splenocytes were isolated from three WT and three *Mtv8*^*KO*^ 10.5-week-old B6J female mice. Red blood cell lysed splenocytes were labeled with microbeads conjugated to monoclonal anti-mouse CD19 antibodies (Miltenyi Biotec, Bergisch Gladbach, Germany) and positively sorted as detailed by the manufacturer. RNA was isolated from sorted cells using guanidine thiocyanate extraction and CsCl gradient centrifugation (*23*).

### Library preparation

Immunoglobulin libraries were generated from 1 μg of RNA using the NEBNext Immune Sequencing Kit (New England Biolabs, Ipswich, Massachusetts, USA) according to manufacturer’s instructions, specifically enriching for B cell receptor (BCR) chains during the first PCR step. The libraries generated from the six individual mice were pooled in equimolar amounts and sequenced by paired-end 300 bp sequencing on an Illumina MiSeq by The University of Chicago Genomics Facility.

### B cell receptor sequence processing and analysis

Preprocessing of BCR sequences was performed using the open-source workflow pRESTO NEBNext Immune Sequencing Kit Workflow (v3.2.0) on Galaxy (*22*). Reads with a Phred quality score <20 were removed for quality control. Reads that did not match to the constant region primer (maximum error rate 0.2) were removed. Reads that did not match to the template switch sequence (maximum error rate 0.5) were removed. The first 17 bp following the template switch site were a unique molecular identifier (UMI) on each read. Sequences with identical UMIs were collapsed into consensus sequences with sequences found in less than 60% of reads removed. Positions with more than 50% gap sequences were removed. Mate-pairs were assembled with a minimum of 8 bp overlap (maximum error rate of 0.3). Assembled reads were assigned isotype-constant region identities based on local alignment of the 3’ ends of the reads (maximum error rate of 0.3). V, (D), and J gene segments were assigned using MiGMAP mapper (Galaxy Version 1.0.3+galaxy2). Subsequent analyses were done using R Studio. Statistically significance was calculated using unpaired Welch’s *t* test with Bonferroni correction. Graphing was done using GraphPad Prism version 10.0.1 for Mac (GraphPad Software, Boston, Massachusetts, USA).

## Data availability

The accession no. for the Ig NEBNext Immune Sequencing datasets reported in this paper is Gene Expression Omnibus no. GSE248410.

## Supporting information

Mtv8_Ig_Repertoire_Summary

## Author contributions

H.A.B. conceived of the project and performed all the experiments reported in this paper. S.A.E. and T.G. designed the CRISPR/Cas9 approach to target *Mtv8*. T.G. maintained the mouse colony. H.A.B wrote the manuscript. T.G. and S.A.E. edited the manuscript.

## Acknowledgments

We thank The University of Chicago Transgenic Mouse Facility (RRID:SCR_019171), especially Linda Degenstein, for the generation of the *Mtv8-*deficient B6J mouse line. We thank The University of Chicago Genomics Facility (RRID:SCR_019196), especially Dr. Pieter Faber, for their assistance with sequencing Ig libraries. We thank Dr. Alexander Chervonsky for helpful discussions. We also thank Dr. David G. Schatz for advice and guidance on the project. H.A.B. was partially supported by the Duchossois Family Institute at the University of Chicago.

## Notes

### Competing Interest Statement

The authors have declared no competing interest.

